# Development of novel lipidic particles for siRNA delivery that are highly effective after 12 months storage

**DOI:** 10.1101/390641

**Authors:** Daniel Clarke, Adi Idris, Nigel AJ McMillan

## Abstract

Liposomes are versatile and well-proven as a means to deliver nucleic acids into cells. Most of the formulation procedures used are labour intensive and result in unstable end products. We have previously reported on the development of a simple, yet efficient, hydration-of-freeze-dried-matrix (HFDM) method to entrap siRNA within lipid particles. Here we show that the particles are stable up to 12 months after storage room temperature (RT), 4°C or - 20°C. While RT storage results in changes in particle size and polydispersity, gene silencing of all particles was similar to freshly prepared particles following storage for 3, 6, 9 or 12 months at all temperatures. This is the first report of such long-term stability in siRNA-loaded liposomes.

## INTRODUCTION

Various methods for formulating polynucleotide-loaded PEGylated particles have been reported to date, including post-insertion ^1^, reverse-phase evaporation ^2^, detergent dialysis ^3^ and ethanol dialysis ^4-6^. However, most of these methods, though effective, require relatively complicated and lengthy formulation procedures with the resulting particles suspended in an aqueous state. This has led to long term storage issues including aggregation and/or fusion of the particles ^7, 8^, hydrolysis of the lipids ^7^, and instability of siRNA nucleotides in an aqueous environment. Moreover, these formulations are also prone to be affected by stresses occurred during transport, such as agitation or temperature fluctuation ^8, 9^. These problems, along with the significantly increased effort required for large-scale production of these particles using the existing formulation procedures has held back clinical development and adoption.

To address these issues, we developed a novel method to formulate stable, siRNA-loaded, PEGylated lipid particles using the hydration of freeze-dried matrix (HFDM) method^10^. This simple, yet-efficient, procedure results in lipid particles that are have highly favourable characteristics for *in vivo* delivery. Indeed, these particles have been used for *in vivo* systemic delivery in animal models to target a range of cancer-related genes, which have resulted in significant tumour elimination and increased survival ^11, 12^. HDFM lipoplexes have also been used to reduce gene expression in lung tissues ^13^, the peritoneal cavity via intraperitoneal delivery ^14^ and in the vaginal epithelia with the aid of a novel aliginate matrix to increase retention time ^15^.

The simple mixing of lipids (DOTAP, cholesterol and PEG2000-C_16_Ceramide – molar ratio of 45:45:10) dissolved in *tert*-butanol with nucleic acids dissolved in sucrose followed by freeze-drying and rehydration results in lipid particles that are isotonic and ready for direct *in vivo* injection. Particles are ~190nm in size with a polydispersity index of 0.32 and an overall charge of 45mV. The encapsulated siRNA was protected from serum ^10^ and show excellent pharmacokinetics with T_1/2_ λz>40 h compared to other systems such as galactosylated cationic liposomes (T_1/2_ λz>1h)^16^, PEGylated polyplexes (T_1/2_ λz>1.5 h)^17^, or SNALPS (T_1/2_ λz>6.5 h)^18^. This represented an advance in lipid formulation of siRNA for *in vivo* use.

As mentioned above, liposomes are prepared fresh for *in vivo* use as they aggregate, fuse or hydrolyse in aqueous solutions over time. As the HFDM method results in a dried matrix we postulated that these resulting lipoplexes would be highly stable over time. Indeed, our initial studies examined the longevity of the freeze-dried siRNA/lipid matrix stored at 4°C, or room temperature, for 4 weeks and showed that silencing was still significant at this time ^10^. Here, we have extended these studies out to 12 months and looked at a range of different storage conditions. We examined the physical characteristics of the particles and their ability to silence target genes *in vitro* at various time points. Our data show that HFDM lipoplexes are highly stable and still active for *in vitro* silencing even 12 months post-production. Such post-production longevity has never previously been reported and represents a significant advance in the field.

## MATERIALS AND METHODS

### Cells and siRNA

HeLa cells were originally obtained from the American Type Culture Collection (ATCC) and were cultured as described previously ^23^. The siRNA used in this study was Lamin A/C siRNA ^24^ obtained from Genesearch (Shanghai, China). siGlo red (Dharmacon, Lafayette, CO) was used as a qualitative transfection indicator as transfection efficiency.

### Liposome formulations

Lipoplexes were prepared by Hydration of Freeze-Dried Matrix (HFDM) method as described previously ^10^. Required amounts of DOTAP, cholesterol and PEG2000-C16Ceramide were dissolved in 1 mL of tert-butanol at a molar ratio of 45:45:10. 40 μg of siRNA was added to 1 mL of filtered sucrose solution before mixing with the lipid solution. The resultant formulation was then snap-frozen and freeze-dried overnight (ALPHA 1–2 LDplus, Martin Christ, Germany) at a condensing temperature of −80°C and pressure of less than 0.1 mbar. Distilled H2O was then added to the lyophilised product with gentle shaking. A Nitrogen/Phosphate (N/P) ratio of 4:1 was used for all formulations and three separate batches were made for each formulation condition (n=3). The final product contained 40 μg siRNA in 300 μL isotonic sucrose solution.

### Particle Characterisation

Size, polydispersity and zeta potential of the resultant lipoplexes were measured using a Zetasizer Nano ZS™ (Malvern Instruments, Malvern, UK) following appropriate dilution in distilled water. Measurements were carried out at room temperature (RT), 4°C and −20°C with 10 runs per measurement undertaken.

### siRNA entrapment efficiency

siRNA entrapment efficiencies were determined using the Quant-iT™ PicoGreen^®^ reagent (Invitrogen) as previously described ^10^.

*Lipoplex cell uptake analysis*

Cells were forward-transfected with 40μM of siGlo red. Cell uptake efficiency was assessed by flow cytometry 24h post-transfection on a BD LSR FORTESSA^TM^ cell analyser (BD bioscience, San Jose, CA)

### In vitro siRNA transfection

*In vitro* siRNA transfection of liposome-complexed siRNA was performed as previously described ^10^. HeLa cells were seeded the day before the transfection experiment at a density of 75,000 cells per well in a 6 well plate. A final concentration of 40 nM liposome-entrapped siRNA suspended in primocin-free complete media was then added to each well. siRNAs were left on cells for 8 h, and cells were then incubated in primocin-containing media for 3 days at 37°C. The level of gene knockdown was determined by qPCR analysis.

### qPCR

Extract of RNA, cDNA generation, and qPCR were carried out as described previously ^23^. Real time primers used: Lamin A/C forward 5’-TGGAGATGATCCCTTGCTGA-3’, Lamin A/C reverse 5’-GCATGGCCACTTCTTCCCA-3’, b-actin forward 5’- AGCCTCGCCTTTGCCGA-3’and b-actin reverse 5’-CTGGTGCCTGGGGCG-3’.

## RESULTS AND DISCUSSION

We have previously showed that storage of our formulated lipoplexes for 1 month at room temperature (RT) and 4°C did not affect its physiochemical properties ^10^. However, the physiochemical stability of the lipoplexes have never been tested outside these storage conditions. The stability of dried-state lipoplexes was investigated up to 12 months post-manufacture. Following formulation, the particles were stored at either RT, 4°C or −20°C. Compared to freshly-prepared lipoplexes, the particle size, zeta potential and polydispersity index (PDI) of the lipoplexes did not vary over time a teach storage condition (Figure 1). Lipoplex powder stored at RT displayed greater variation in both size and PDI upon rehydration, likely due to varying humidity levels during storage (Figure 1A, Figure 1B). More importantly, no variation in the surface charge (or zeta potential) of particles was observed when stored at any temperature over a 12-month period (Figure 1C), which is important for the stability of the particles and remaining free from agglutination when delivered into the bloodstream. In contrast to what we previously observed ^10^, the siRNA entrapment efficiency did not fluctuate over time at different temperatures when compared to freshly-prepared lipoplexes, indicating the siRNA encapsulated in the lipoplexes were stable (Figure 1D). Overall, there was minimal effect of the storage time and temperature on the lipoplex physiochemical properties and the stability of the siRNA-loaded lipoplexes, with 4°C and −20°C showing superior stability in terms of size, charge and size variation.

**Figure 1.**
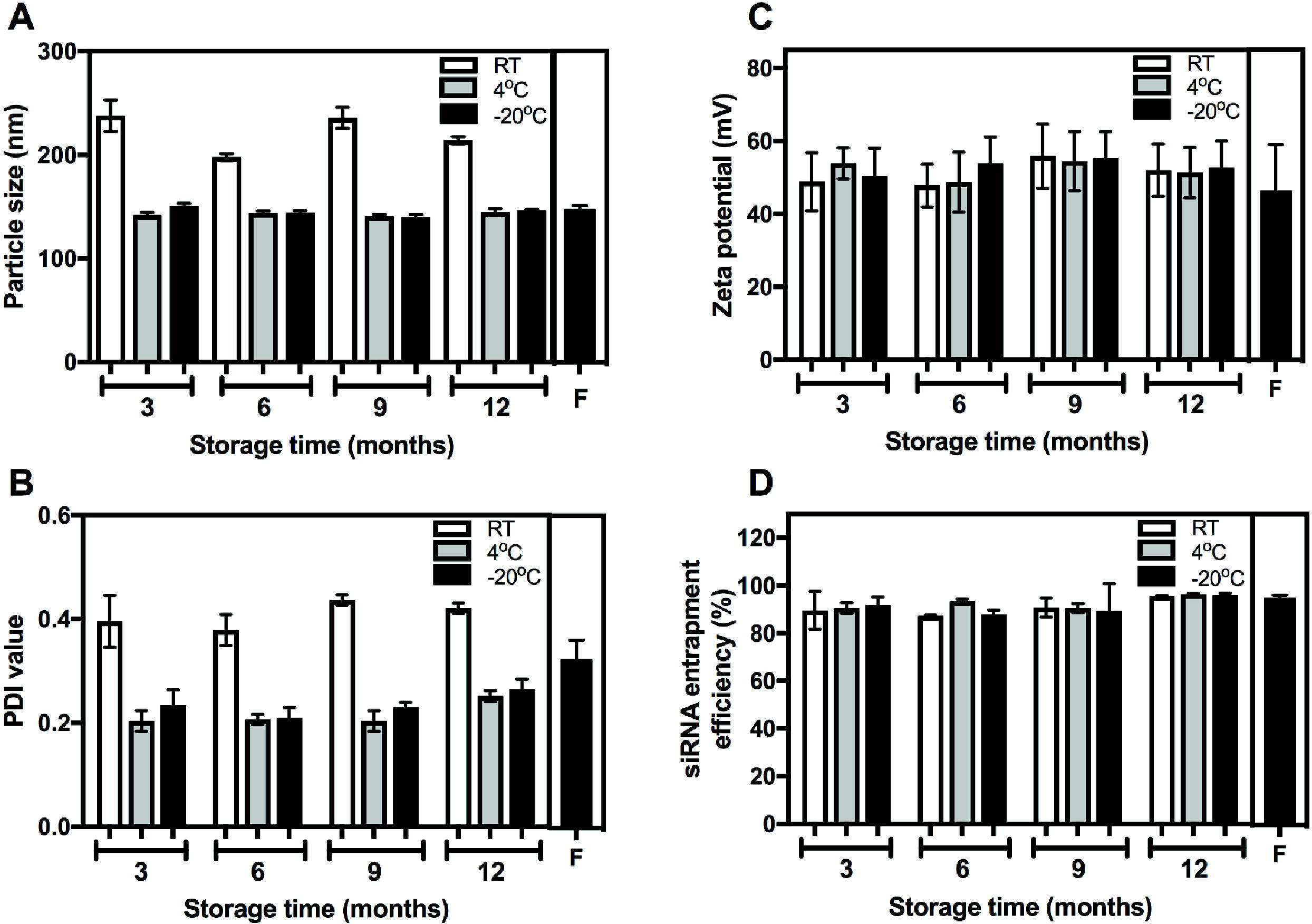
Lipoplex size, polydispersity, zeta potential and siRNA entrapment efficiency remains unchanged with long term storage at various temperatures. (A) Size, (B) polydispersity and (C) zeta potential of the resultant lipoplex were measured using a Zetasizer Nano ZS following appropriate dilution in distilled water. Formulations were stored at either −20°C, 4°C or room temperature (RT) and 2 separate batches were made for each condition. (n=2). (D) siRNA entrapment efficiencies were determined using the Quant-iT™ PicoGreen^®^ reagent. Tests were performed on freshly prepared lipoplex (F – Black shaded bars), and lipoplex stored at various temperatures for different lengths of time post manufacture. Bar graphs represent the mean and the error bars represent the standard deviation.

To assess whether storage time and temperature have any effect on the functional capability of the lipoplexes, the gene knockdown ability of siRNA-loaded lipoplexes was tested *in vitro*. *In vitro* cell uptake efficiency of lipoplexes stored over time at varying temperatures were comparable to that of freshly prepared lipoplexes (Figure 2A). Liposome powders were rehydrated at each time point, complexed with siRNAs and applied to HeLa cells to determine knockdown of the non-essential structural protein, lamin A/C. qPCR analysis revealed over 50% decrease of lamin A/C mRNA expression in HeLa cells treated with A/C-specific siRNA compared to untransfected cells, irrespective of the storage conditions (Figure 2B). The degree of gene knockdown was comparable to that of freshly prepared liposomes suggesting that storage time and temperature have minimal effect on the gene-silencing efficiency of the siRNAs-complexed with lipoplexes.

**Figure 2.**
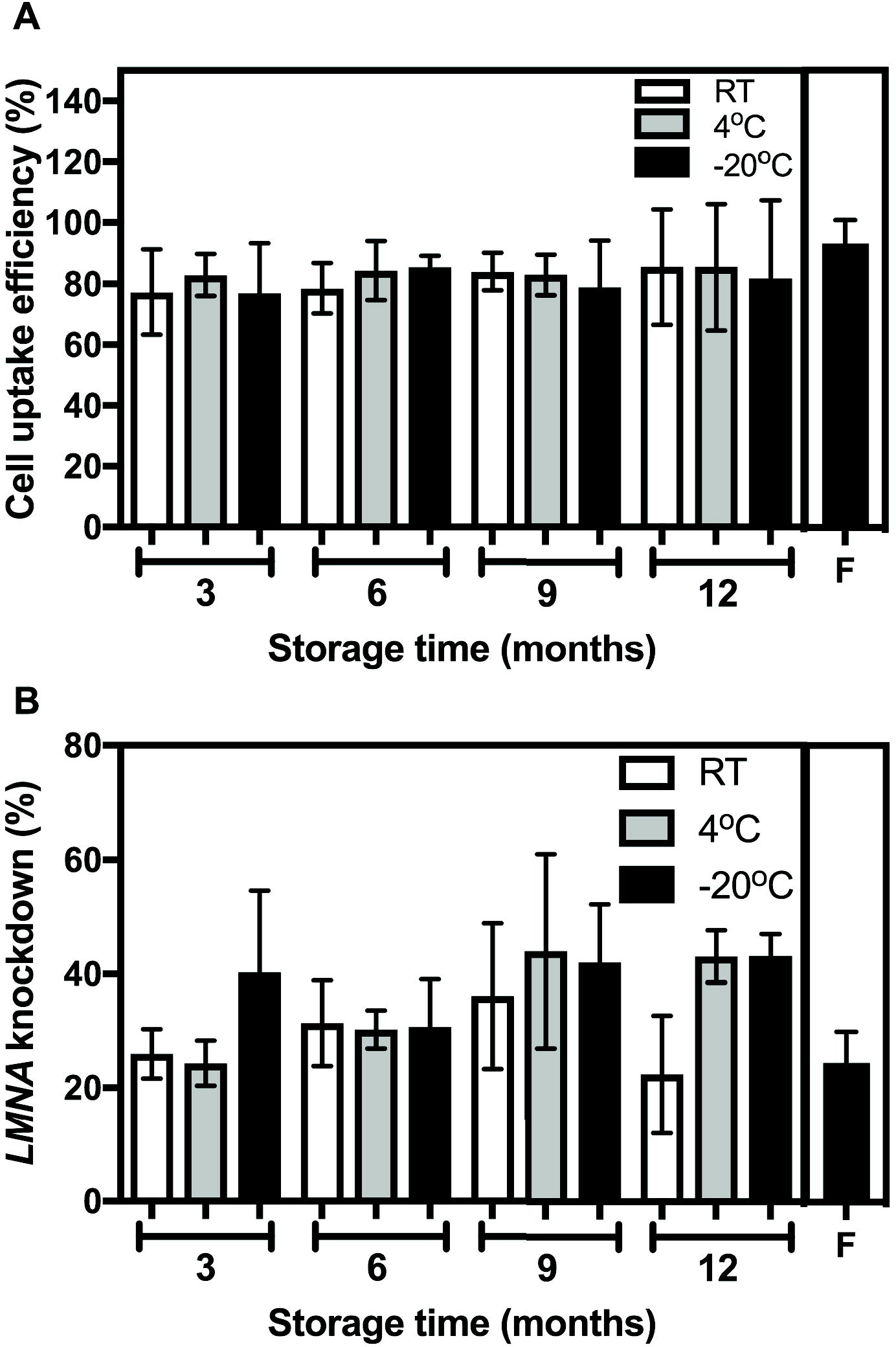
siRNA-mediated target gene knockdown capability was maintained using lipoplexes stored at different temperatures over time *in vitro*. A) HeLa cells were treated with a final concentration of 40 nM of siGlo complexed into lipoplexes and incubated at 37°C for 24h before assessing siGlo uptake (%) by flow cytometry. Tests were performed on freshly prepared lipoplexes (F – Black shaded bars), and lipoplexes stored at various temperatures for different lengths of time post manufacture. Bar graphs represent the mean and the error bars represent the standard deviation from three independent experiments. B) HeLa cells were treated with a final concentration of 40 nM of Lamin A/C siRNA complexed into lipoplexes and incubated at 37°C for 3 days. Expression levels were calculated relative to non-transfected cells by qPCR analysis. Tests were performed on freshly prepared lipoplexes (F – Black shaded bars), and lipoplexes stored at various temperatures for different lengths of time post manufacture. Bar graphs represent the mean and the error bars represent the standard deviation from three independent experiment.

Previous studies have investigated the stability of dried-state liposomal preparations at different temperatures (i.e. 37°C to 60°C) over a 1-month period ^19^, this is the first study to investigate the physiochemical and functional effects of the prolonged storage of lipoplexes over a longer period of time (i.e. 12 months) and at lower temperatures. We developed this lyophilization procedure to ensure effective long-term storage of our lipoplexes, even at room temperature, as lipoplex suspensions are known to be unstable in aqueous suspension for long-term storage, especially with respect to hydrolysis and size stability. To date no data on the long term (up to 12 months) viability of siRNA-loaded liposomes have, to our knowledge, been published. There are longevity studies investigating traditional liposomes loaded with plasmid DNA. For example, an early study of unextruded liposomes made from DMRIE:DOPE complexed with plasmid DNA showed that when frozen as a aqueous solution they retained activity when thawed 12 months later ^20^. More recently it was showed that DC-Cholesterol: DOPE liposomes loaded with plasmid DNA and freeze-dried were stable following 3 months storage at RT and 40°C, although storage at 60°C resulted in lost DNA structure ^21^. Others have investigated the stability of DNA oligonucleotide-loaded liposomes for antisense therapies, which changed size and lost ODN content over 12 months (RT) or 90 days (frozen)^22^. However, the functionality of these stored lipoplexes was not examined. Overall, we show that RT was inferior for storage, resulting in an increased particle size and polydispersity from 3 months onwards. Storage at 4°C and −20°C resulted in no changes in lipoplexes and silencing ability. Overall, HFDM liposomes exhibit excellent long-term storage characteristics.

## Acknowledgements

The authors would like to acknowledge funding from the National Health and Medical Research Council of Australia, the Cancer Council Queensland and the Menzies Health Institute Queensland.

